# Targeting endogenous K-RAS for degradation through the affinity-directed protein missile system

**DOI:** 10.1101/805150

**Authors:** Sascha Röth, Thomas J. Macartney, Agnieszka Konopacka, Markus A. Queisser, Gopal P. Sapkota

## Abstract

For over three decades, *K-RAS* has been known as the holy grail of cancer targets, one of the most frequently mutated oncogenes in cancer. Because the development of conventional small molecule K-RAS inhibitors has been extremely challenging, K-RAS has been dubbed as an undruggable target, and only recently a mutation specific inhibitor has reached clinical trials. Targeted protein degradation has emerged as a new modality in drug discovery to tackle undruggable targets. However, no degrader for K-RAS has been described thus far. Our laboratory has developed an Affinity-directed PROtein Missile (AdPROM) system for targeted proteolysis of endogenous proteins through the ubiquitin proteasome system. Here, we show that we can achieve degradation of endogenous K-RAS and H-RAS in different cell lines in a targeted manner using our AdPROM system. Our findings imply that endogenous RAS proteins can be targeted for proteolysis, thereby offering tantalising possibilities for an alternative therapeutic approach to these so-called undruggable targets in cancer.

## Background

The three RAS oncogenes *H-RAS, K-RAS* and *N-RAS*, represent the most frequently mutated genes in cancer [1, 2]. They encode four highly similar proteins, namely H-RAS, N-RAS, K-RAS4A and K-RAS4B, which undergo C-terminal farnesylation [3, 4]. Farnesylation, in combination with palmitoylation in the hypervariable region (HVR) (N-RAS, H-RAS, K-RAS4A) or with a polybasic signal in the HVR (K-RAS4B), mediates the plasma membrane interaction [5]. RAS proteins are small GTPases, which cycle between the GTP-bound (active) and GDP-bound (inactive) states, controlled by guanosine nucleotide exchange factors (GEF) and GTPase activating proteins (GAPs) [6]. Activation of RAS proteins by various extracellular growth factors initiates activation of numerous downstream signalling networks, including BRAF/MAPK and PI3K pathways [7], that are critical for cell proliferation and viability. Many pathogenic mutations in *RAS* genes impair GAP mediated GTP hydrolysis, thereby favouring the persistence of the active RAS-GTP state, which triggers constitutive activation of downstream signalling resulting in unchecked proliferation of cancer cells [2, 8].

As the oncogenicity of RAS mutations has been known for over three decades, intensive efforts have been made towards drugging them. These efforts are yet to result in effective RAS-inhibitor therapies [1, 9]. This has promoted the perception that RAS proteins are undruggable. Several factors make RAS proteins difficult targets to engineer selective small molecule inhibitors. First, the relatively high concentrations of GTP and GDP in cells and picomolar affinity to binding RAS proteins makes it almost impossible to develop GTP/GDP analogues as inhibitors [1, 10]. Second, structural analysis of RAS proteins revealed few sufficiently large and deep hydrophobic pockets on the surface for small molecule binding [11, 12]. Recently, a covalent inhibitor targeting a cysteine in K-RAS G12C was developed to target this specific mutation [13]. However, these barriers and failure to directly target RAS have prompted researchers to explore targeting upstream regulators, or downstream effectors of RAS proteins [1,9,14–16], as well as altering levels of RAS protein, for example by inducing targeted degradation of RAS [17].

Most targeted protein degradation approaches harness the cellular proteolytic pathways that naturally maintain proteostasis, with the ubiquitin-proteasome system (UPS) being frequently exploited [18]. Protein degradation by the UPS is triggered by conjugation of ubiquitin chains onto the target protein, which is achieved through a sequential action of three enzymes: the ubiquitin-activating enzyme (E1), which activates the carboxy-terminal glycine residue of ubiquitin in an ATP-dependent manner; a ubiquitin-conjugating enzyme (E2), which conjugates the activated ubiquitin to its active site cysteine; and a ubiquitin ligase (E3), which facilitates the transfer of ubiquitin from E2 to primarily lysine residues on substrate proteins [19, 20]. Further ubiquitylation on one or more lysine residues within ubiquitin then triggers polyubiquitylation, followed by degradation by the proteasome [21–23]. Targeting RAS for proteolysis relies on the engagement of the cellular proteolytic systems for its ubiquitylation and degradation. In this context, it has been shown that the heterobifunctional molecule dTAG-13, which recruits FKBP12_F36V_-tagged proteins of interest (POIs) to the CRBN/CUL4A E3 ubiquitin ligase for their degradation, can degrade FKBP12_F36V_-KRAS_G12V_ overexpressed in cell lines [17]. However, FKBP12_F36V_ itself can be targeted for ubiquitylation when using heterobifunctional small molecule binders [24]. Therefore, it remains unclear, whether using dTAG13 on FKBP12_F36V_-K-RAS results in the ubiquitination of K-RAS or FKBP12_F36V_. Such information is not only key to evaluate proteolysis as a druggable approach for targeting RAS proteins but also to inform on the development of effective heterobifunctional RAS degraders.

We have previously developed an effective proteolytic Affinity-directed PROtein Missile (AdPROM) system for UPS mediated POI degradation [25, 26]. AdPROM consists of a fusion of von-Hippel-Lindau (VHL) protein, a substrate recruiter of the CUL2-RING E3 ligase complex, and high-affinity binders, such as nanobodies and monobodies, of POIs. Delivering AdPROM into multiple cell lines through retroviral transductions led to efficient degradation of endogenous target proteins, including SHP2 and ASC [26]. Furthermore, in order to target POIs for which no high-affinity polypeptide binders exist, we utilized CRISPR/Cas9 genome editing to rapidly introduce GFP tags on endogenous VPS34 and PAWS1 genes, and used the AdPROM system consisting of anti-GFP nanobody fused to VHL to achieve near-complete degradation of the endogenous GFP-VPS34 and PAWS1-GFP proteins [25]. In this study, we explore the use of the AdPROM system, and demonstrate its efficacy, for targeted degradation of endogenously GFP-tagged K-RAS and untagged, endogenous K-RAS from cells.

## Methods

### Sequence Alignment

Protein sequences of K-RAS4A/B, H-RAS and N-RAS were taken from Uniprot [27] and aligned in Clustal Omega [28]. The alignment was further processed in JalView [29] to highlight percent sequence identity.

### RNA extraction, cDNA synthesis and qRT-PCR

For RNA extraction, 2×10_5_ cells were seeded in a 6-well dish and harvested the next day with the RNeasy Micro Kit (Qiagen, #74004) according to the manufacturer’s protocol. 1 μg of RNA was reverse transcribed with the iScript cDNA synthesis Kit (BIORAD, #1708891) according to the manufacturer’s protocol. For qRT-PCR 1 μl of diluted cDNA (1:20 or 1:80) was mixed with forward and reverse primers (Custom primers from Invitrogen, 300 nm final concentration each) and SsoFast EvaGreen Supermix (BIORAD, #1725204) in a 384-well plate (Axygen, #321-22-051) and run on a BIORAD CFX384.

Primer sequences:

K-RAS4A fw: GAGGGAGATCCGACAATACAG;

K-RAS4A rev: TCTCGAACTAATGTATAGAAGGCATC;

K-RAS4Bfw: TTGCCTTCTAGAACAGTAGACAC;

K-RAS4B rev: CATCGTCAACACCCTGTCTTG;

Total K-RAS fw: GGAGTACAGTGCAATGAGGG;

Total K-RAS rev: CCATAGGTACATCTTCAGAGTCC;

H-RAS fw: GAACAAGTGTGACCTGGCT;

H-RAS rev: ACCAACGTGTAGAAGGCATC;

N-RAS fw: AATACATGAGGACAGGCGAAG;

N-RAS rev: GTTTCCCACTAGCACCATAGG;

GAPDH fw: CTTTGTCAAGCTCATTTCCTGG;

GAPDH rev: TCTTCCTCTTGTGCTCTTGC.

Melting curves were analysed for purity of the PCR product and fold changes were calculated by the 2-ΔΔCt method [30].

### Cell line maintenance and manipulation

All cells were cultured in humidified incubators at 37°C and 5% CO_2_. A549, HEK293-FT, A375, A172 and SW620 cells were cultured in Dulbecco’s modified Eagle’s medium (DMEM; Gibco) with 10% FBS (Sigma), 1% penicillin/streptomycin (Lonza) and 2 mM L-glutamine (Lonza). HT-29, HPAFII and H460 cells were cultured in RPMI1640 medium (Gibco), with the same supplements as DMEM. For retrovirus production, 3.2 μg pCMV-gag-pol, 2.2 μg pCMV-VSV-G and 6 μg of respective pBabeD plasmids were co-transfected in roughly 70% confluent HEK293-FT cells cultured on a 10-cm dish. Plasmids were mixed with 600 μl Opti-MEM (Gibco) and 24 μl of 1 mg/ml polyethyleneimine (Polysciences) dissolved in 25 mM HEPES pH 7.5. The mixture was vigorously vortexed for 15 s and incubated for 20 min at room temperature. The volume was adjusted to 10 ml with DMEM and added to FT cells. After 24 h, medium was exchanged to DMEM or RPMI, depending on the target cell growth medium. After an additional 24 h, the medium was harvested and filtered through a 0.45 μm Minisart syringe filter (Sartorius). The supernatant was added to a plate of roughly 70% confluent target cells in a 1:10–1:4 dilution (in respective medium) in the presence of 8 μg/ml polybrene (Sigma). After 24 h, growth medium was exchanged with fresh medium containing 2 μg/ml puromycin, to select transduced cells. Puromycin was removed from the medium after 48 h.

Cells were lysed on ice, by washing once with PBS and scraping in lysis buffer (50 mM Tris–HCl pH 7.5, 0.27 M sucrose, 150 mM NaCl, 1 mM EGTA, 1 mM EDTA, 1 mM sodium orthovanadate, 1 mM sodium β-glycerophosphate, 50 mM sodium fluoride, 5 mM sodium pyrophosphate, 1% (v/v) Triton X-100 and 0.5% Nonidet P-40) supplemented with protease inhibitors (Roche; 1 tablet/25 ml of lysis buffer). Protein content from cleared cell lysates was determined with Pierce Detergent Compatible Bradford Assay Kit (Thermo Fisher). Lysates were processed further or frozen and stored at -20°C.

### CRISPR/Cas9

For generation of N-terminal GFP knock-in A549 cell lines the K-RAS locus was targeted with a dual guide approach [31] (using the sense guide (pBabeD vector, DU54976): GCGAATATGATCCAACAATAG; antisense guide (pX335 vector, DU54980): GCTGAATTAGCTGTATCGTCA; and the GFP-KRAS donor (pMK-RQ vector, DU57406). Briefly, 1 μg of each of the guideRNA plasmids and 3 μg of the donor plasmid were co-transfected into A549 cells. Plasmids were mixed with 1 ml of Opti-MEM (Gibco) and 20 μl of 1 mg/ml polyethyleneimine (Polysciences), vortexed vigorously for 15 s and added to 70% confluent cells in a 10-cm dish. The next day, cells were selected in puromycin (2.5 μg/ml) for 48 h and re-transfected with the same plasmids once they reached 70% confluence. Single GFP positive cells were obtained by FACS sorting and surviving single cell clones were screened by genomic DNA based PCR and western blot to validate homozygous knockin of the GFP-tag on the endogenous *KRAS* gene. For PCR based screening the following primers were used: Fw: ATCCAAGAGAACTACTGCCATGATGC; Rv: CATGACCTTCAAGGTGTCTTACAGGTC. PCR products of positive clones were cloned with the StrataClone PCR Cloning Kit (Agilent) into the supplied vector system, according to the manufacturer’s protocol. Sequencing of positive clones was carried out by the MRC-PPU DNA Sequencing and Services with a custom primer close to the RAS mutation site (Rv: CAAAGAATGGTCCTGCACCAG).

### SDS PAGE and Western Blotting

Cell lysates were adjusted to uniform protein concentration and mixed with 6x reducing Laemmli SDS sample buffer (Fisher Scientific). 10-20 μg of total lysate protein, or immunoprecipitates were resolved by SDS polyacrylamide gel electrophoeresis (PAGE). After PAGE, proteins were transferred onto methanol activated PVDF membrane (Immobilon-P or Immobilon-FL, Merck) in Tris/glycine buffer containing 20% methanol in a tank blotting system for 85 min at a constant voltage of 85 V. The membranes were then re-incubated with methanol for 2 minutes and stained with Ponceau S solution to gauge uniform protein transfer (Sigma). After de-staining membranes in TBS-T (50 mM Tris–HCl pH 7.5, 150 mM NaCl, 0.1% Tween-20), they were blocked for 1 h in 5% non-fat milk (Marvel) in TBS-T. Primary antibody incubation was done overnight at 4°C in 5% milk/TBS-T. Following 3×10 min washes in TBS-T, membranes were incubated with respective HRP-conjugated (CST) or fluorescently labelled (Biorad) secondary antibodies for 1 h, washed again 3×10 min in TBS-T and developed on a ChemiDoc gel imaging system (Biorad) using the respective channels. HRP-conjugated blots were incubated with Immobilon Western Chemiluminescent HRP Substrate (Millipore).

### Immunoprecipitation

Cell lysates were adjusted to 1 μg/μl in lysis buffer. Either GFP-trap beads (ChromoTek) or Anti-FLAG-M2-Affinity agarose resin (SigmaAldrich) was equilibrated with lysis buffer. 300-500 μg of total protein was added to 10-15 μl of beads (50% slurry) and incubated for an hour at 4°C under agitation. Centrifugation steps at 200xg were done at 4°C for 2 minutes. Supernatant (flowthrough) was separated from beads, and beads were washed 3-5 times in lysis buffer. Proteins were eluted in lysis buffer containing Laemmli SDS sample buffer by boiling at 95°C for 5 minutes.

### Antibodies

Antibodies were purchased from Thermo Fisher (Alpha tubulin, MA1-80189; rat-HRP, 31470), Abcam (panRAS, ab206969; HIF1a, ab1), Sigma (K-RAS4B, WH0003845M1; Flag-HRP, A8592-.2MG; GFP, 11814460001) CST (GAPDH, 2118S; rabbit-HRP, 7074S; mouse-HRP, 7076S) and Bio-Rad (rabbit starbright 700, 12004161). Primary antibodies were generally used in 1:1,000 dilutions in 5% milk TBS-T, apart from RAS (1:500), and GAPDH & alpha-tubulin (1:5,000). Secondary antibodies were used in a 1:5,000 dilution in 5% milk TBS-T. Other primary antibodies recognizing different RAS species were obtained from Proteintech (N-RAS, 10724-1-AP; H-RAS, 18295-1-AP; K-RAS2B, 16155-1-AP; K-RAS2A, 16156-1-AP) and Invitrogen (H-RAS, PA5-22392; K-RAS, 415700).

Antibodies for immunofluorescence were purchased from MBL/Caltag Medsystems (GFP, 598), Abcam (ATPB, ab14730), BD Biosciences (P120 Catenin, 610133), Sigma (Flag-M2, F1804) and Thermo Fisher (AlexaFluor488 [donkey anti-rabbit], A21206; AlexaFluor594 [goat anti-mouse], A11005).

### Immunofluorescence

Cells were seeded in a 12-well dish onto cover slips and grown over night. The next day, cells were washed twice in PBS and fixed for 10 minutes in 4% formaldehyde/PBS (Sigma). Coverslips were washed in DMEM (Gibco) containing 10 mM HEPES followed by a 10 min incubation. Coverslips were washed in PBS and permeabilised for 3 min in either 0.2% NP-40/PBS or 0.2% Triton X-100/PBS. Coverslips were washed twice in PBS and blocked for 15 min in 3% BSA (Sigma) in PBS. Primary antibody incubation was done for 1-2 h at room temperature at appropriate antibody dilutions in blocking solution. Residual antibody was washed away in 0.2% Tween/PBS (3×10 min). Secondary antibody incubation was done for 30 min at 1:300 antibody dilution in the dark. The same wash steps were repeated, but the first wash contained DAPI (0.5–1 μg in 10 ml). Finally, coverslips were dipped in water, air dried and mounted on slides with Vectashield (Vector Laboratories). Fluorescence signals were analysed on a Deltavision Widefield microscope (GE). Images were deconvolved using the default settings of softWoRx Imaging software.

### Cell Proliferation Assays

After trypsinization, live cell numbers were determined in a Neubauer haemocytometer in the presence of trypan blue. Cell numbers were adjusted to 5000 cells per ml in the respective growth medium. 5000 cells were added per well of a 12-well dish, and each line was grown in triplicates. After 7 days, relative cell numbers were determined by crystal violet staining. In short, cells were washed in PBS, fixed for 5 min in fixing buffer (10% methanol, 10% acetic acid), washed in PBS again and incubated for 30-60 min in crystal violet solution (0.5% crystal violet in 20% methanol). Plates were dipped in tap water to remove stain and air dried overnight. Plates were scanned on a Licor Odyssey using the 700 nm channel. Subsequently, 1 ml methanol was added to each well and plates were incubated shaking for 30 min. Depending on the colour of 1 set of cells, 100-200 μl of supernatant was loaded in triplicate on a 96-well plate and absorbance at 570 nm was measured in an Epoch microplate spectrophotometer (BioTek). Values were normalized to the untreated sample and a one-way ANOVA analysis with Dunnett’s multiple comparisons test was done.

### Flow Cytometric Analysis

Cells were trypsinized, washed and resuspended in PBS containing 1% FBS. Cells were then analysed on a FACS Canto II flow cytometer. Cells were analysed with the following gating strategy: (i) cells: in a plot of FSC-A vs. SSC-A, a gate was drawn surrounding the major population of cells, removing debris and dead cells. (ii) single cells: in a plot of FSC-A vs. FSC-W, a gate was drawn around an area corresponding to single cells. (iii) in the ‘single cells’ population on a GFP-A vs. PE-A plot a gate was drawn around GFP-positive cells in A549_GFPKRAS_ sample, using WT A549 cells as a negative control. Gates (i) and (ii) were adjusted to the individual cell lines. Gate (iii) was kept unchanged within an experiment.

## Results

### Generation of a GFP-KRAS knock-in non-small cell lung cancer A549 cell line

The high degree of amino acid sequence similarity between the four RAS proteins, i.e. K-RAS4A, K-RAS4B, H-RAS and N-RAS (Fig. 1A), and the subsequent difficulty in generating selective antibodies against individual isoforms pose substantial challenges in studying specific RAS proteins [32]. In order to explore targeted proteolysis of K-RAS using the AdPROM system, we employed CRISPR/Cas9 technology to generate an A549 non-small cell lung carcinoma (NSCLC) cell line harbouring a homozygous knock-in of green fluorescent protein (GFP) cDNA at the N-terminus of the native *K-RAS* gene (Fig. S1). As K-RAS4A and K-RAS4B are splice variants differing only in their extreme C-terminus (Fig. 1A), this approach allowed us to simultaneously tag both isoforms with GFP. The homozygous GFP knock-ins on the native K-RAS locus (A549_GFPKRAS_) were verified by genomic sequencing (Fig. S1). Moreover, by western blot analysis using both pan-RAS and K-RAS4B antibodies, the appearance of higher molecular weight GFP-K-RAS species with a concurrent disappearance of the native molecular weight K-RAS species was evident in the A549_GFPKRAS_ cell line compared to wild type (WT) A549 control cells (Fig. 1B). The use of a panRAS antibody resulted in the detection of two distinct bands in A549 WT cells (Fig.1B). As the lower band remained intact in A549_GFPKRAS_ cells, it most likely corresponds to H- and/or N-RAS (Fig. 1B). However, in A549 cells we were unable to detect any endogenous signals with commercially available H-RAS, N-RAS or K-RAS4A specific antibodies (listed in Methods section). By qRT-PCR, we showed that levels of H- and N-RAS transcripts were slightly reduced in A549_GFPKRAS_ cells compared to WT A549 cells, while transcript levels of K-RAS were reduced by roughly 50% (Fig. S2). We were able to efficiently immunoprecipitate GFP-K-RAS from A549_GFPKRAS_ but not WT A549 cell extracts (Fig. 1C).

**Figure 1.**
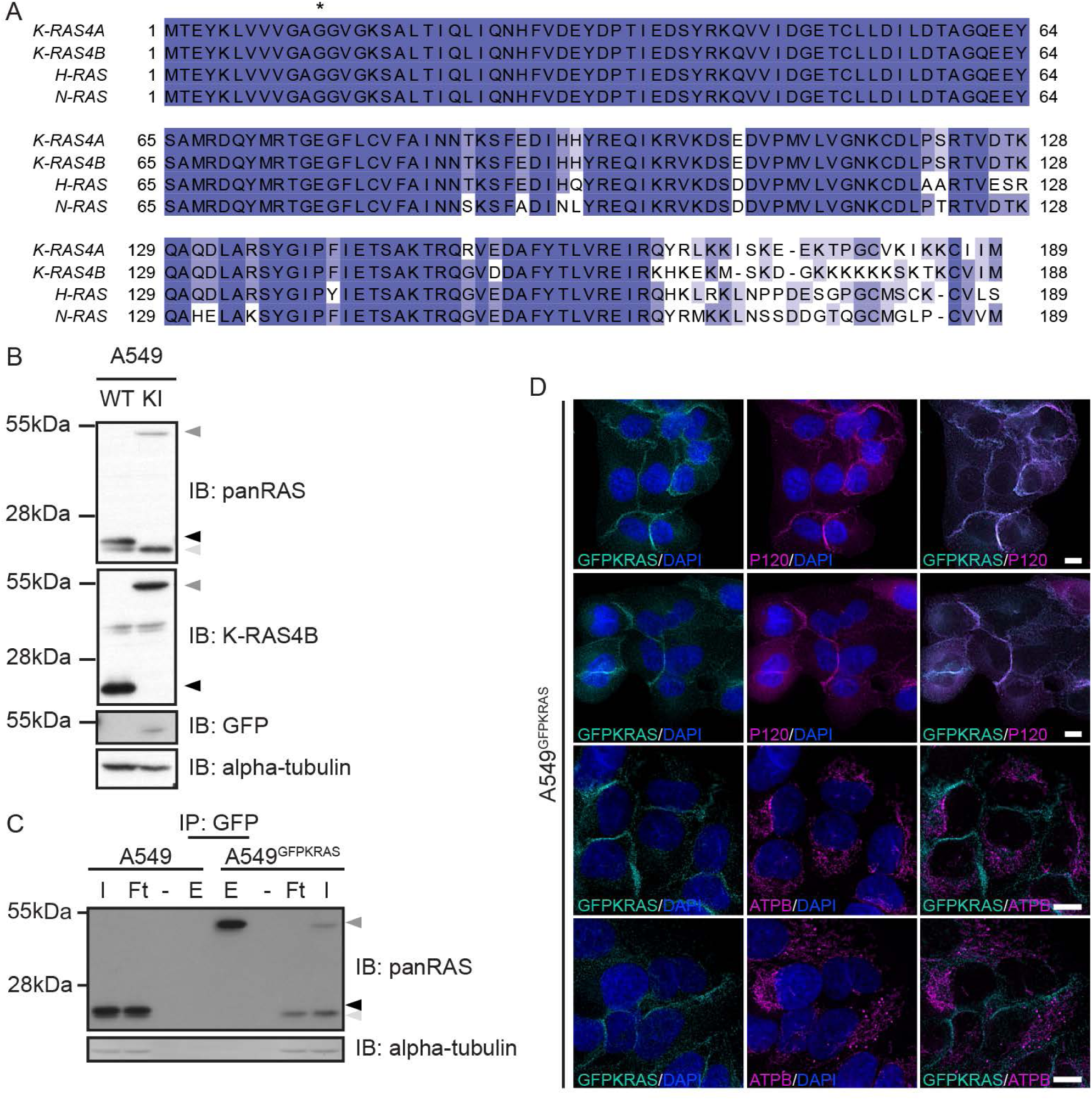
Generation of GFP-KRAS knockin in A549 NSCLC cells by CRISPR/Cas9. (A) Sequence Alignment of RAS protein isoforms K-RAS4A (Uniprot-ID: P01116-1), K-RAS4B (P01116-2), H-RAS (P01112-1) and N-RAS (P01111-1). Degrees of shading according to % sequence identity between the four proteins. Asterisk denotes frequently mutated G12 position. (B) A549 WT or K-RAS_GFP/GFP_ knock-in (KI; hereafter called A549_GFPKRAS_) cell lysates were separated by SDS PAGE and the indicated antibodies were used for detection by Western blotting. Arrows indicate different RAS species (black: endogenous K-RAS; dark grey: GFP-K-RAS; light grey: H-/N-RAS). (C) Lysates were processed as in (B) and subjected to immunoprecipitation with GFP-trap beads. I = Input, Ft = Flowthrough, E = Elution. (D) Widefield immunofluorescence microscopy of untreated A549_GFPKRAS_ cells labelled with antibodies specific for GFP (all left panels, cyan) and P120 (top two middle panels, magenta) or ATPB (bottom two middle panels, magenta), and DAPI (all left and middle panels, blue). Overlay of GFP and P120/ATPB is shown on the right. Scalebar = 10μm. Two representative images for each staining are shown. All blots are representative of at least 3 independent experiments.

Recently, a number of RAS antibodies have been evaluated for selective recognition of the different RAS proteins by Western blotting [32], but none of these have been selective for use in immunofluorescence studies. Consequently, studies evaluating subcellular distribution of RAS proteins have been restricted to overexpression systems. Validation of A549_GFPKRAS_ cells allowed us to investigate the sub-cellular distribution of endogenous GFP-K-RAS driven by the native promoter. Endogenous GFP-K-RAS displayed predominantly plasma membrane distribution, which was confirmed by co-staining with P120 catenin, which is known to localise to the plasma membrane [33] (Fig. 1D, Fig. S3). Additionally, we also observed some weak cytoplasmic localisation of GFP-K-RAS. However, no co-localisation of GFP-K-RAS was observed with mitochondrial marker ATPB [34] (Fig. 1D, Fig. S3).

### Targeted degradation of GFP-K-RAS by the proteolytic AdPROM system

We sought to test whether endogenously expressed GFP-K-RAS protein in A549_GFPKRAS_ cells could be targeted for degradation by AdPROM [25, 26]. We have previously shown that fusion of VHL to an aGFP16 nanobody recruits GFP-tagged proteins, such as VPS34 and PAWS1, to the CUL2-RBX1 E3 ligase machinery for target ubiquitination and subsequent proteasomal degradation [25]. Therefore, we postulated that GFP-K-RAS could be recruited in a similar manner to the CUL2-RBX complex for ubiquitination and degradation (Fig. 2A). Indeed, expression of VHL-aGFP16 AdPROM resulted in near complete clearance of GFP-K-RAS from A549_GFPKRAS_ cells compared to the untransduced controls, while the low molecular weight band corresponding to H- and/or N-RAS was unaffected (Fig. 2B). In contrast, neither VHL nor the aGFP16 nanobody alone, serving as controls, caused any apparent changes in the steady state levels of GFP-K-RAS or other RAS proteins (Fig. 2B). Treatment of VHL-aGFP16 AdPROM expressing A549_GFPKRAS_ cells with the Cullin neddylation inhibitor MLN4924 partially rescued the degradation of GFP-K-RAS compared to DMSO-treated controls (Fig. 2C). The neddylation of CUL2 allows a conformational change of the CUL2-RBX E3 ligase machinery so that the RBX E3 ligase is able to ubiquitinate substrates recruited by VHL. In line with this notion, the levels of HIF1α protein, a *bona fide* substrate of VHL [35], were stabilized upon MLN4924 treatment compared to DMSO control (Fig. 2C). Despite the high apparent efficiency of GFP-KRAS degradation by VHL-aGFP16 AdPROM, the retroviral transduction of A549_GFPKRAS_ cells often generates uneven levels of AdPROM expression in a mixed population of cells. Therefore, in order to get a better understanding of the distribution of the cells within this population, we employed a flow cytometric analysis based on GFP fluorescence. We employed gates to define a GFP-positive population based on the GFP-signal from untransduced A549_GFPKRAS_ cells and using WT A549 cells as a GFP-negative control (Fig. 2D). In accordance with the Western blot results (Fig. 2B), 97% of cells expressing VHL-aGFP16 AdPROM showed GFP-KRAS degradation compared to untransduced A549_GFPKRAS_ cells (Fig. 2D), which manifested in an overall reduction of GFP fluorescence of the single cell population (Fig. 2E). The remaining 3% of A549_GFPKRAS_ cells produced GFP signal comparable to untransduced GFP-positive-population, which could be due to low level AdPROM expression within these cells (Fig. 2D). In contrast, A549_GFPKRAS_ cells expressing VHL or aGFP16 alone were defined as GFP-positive at 99.3% or 99.8%, respectively (Fig. 2D, E).

**Figure 2.**
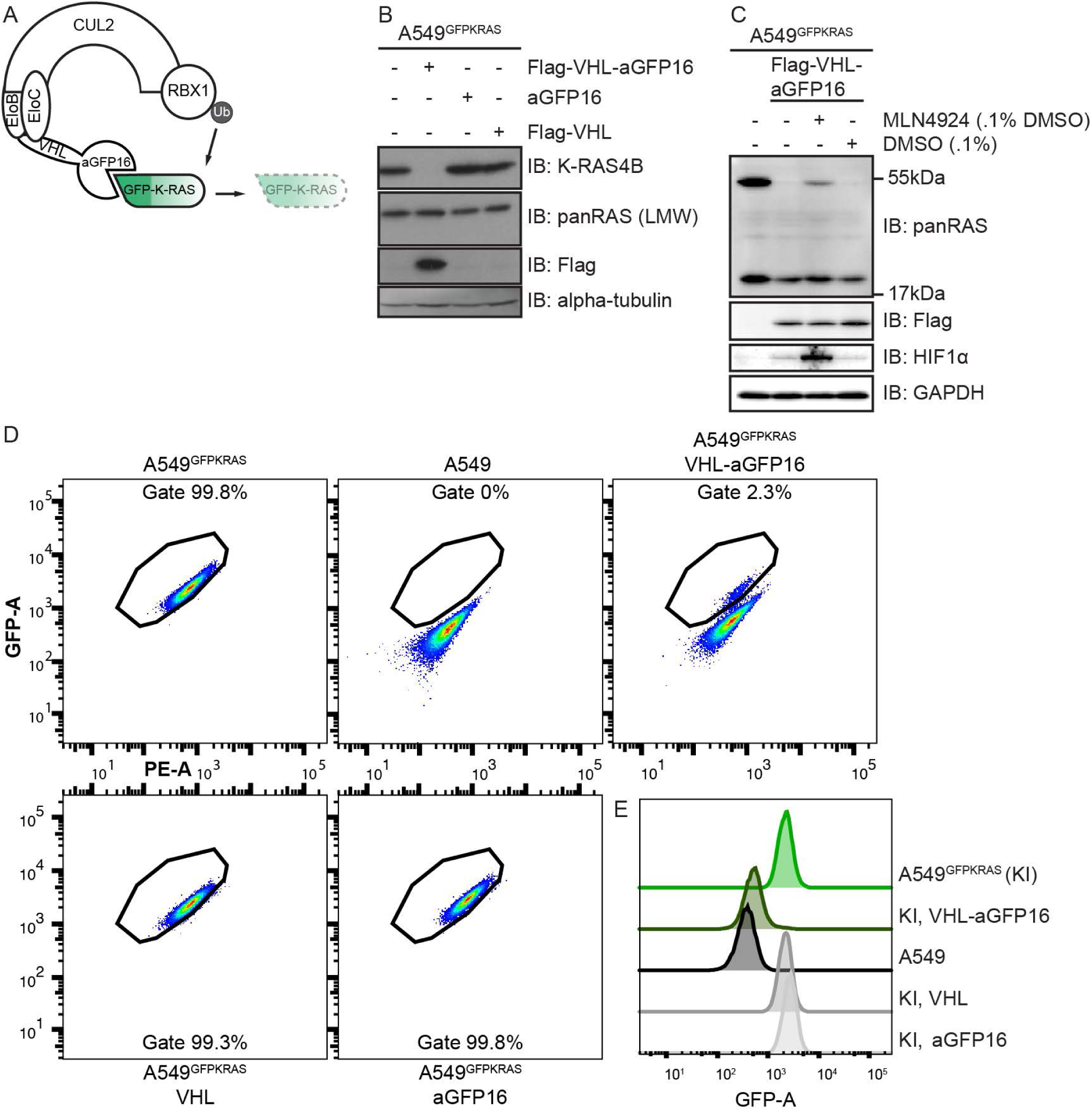
AdPROM mediated degradation of GFP-K-RAS. (A) Schematic representation of the proteolytic AdPROM system. The high affinity GFP-binder aGFP16 is fused to VHL, which is recruited by EloB and EloC to Cul2. aGFP16 recruits GFP-tagged K-RAS and presents it in close proximity to RBX1 in the assembled Cul2 complex. Ubiquitin (Ub) is transferred onto K-RAS, which is subsequently degraded (dashed lines and faded). (B) After treatment with retroviruses and selection, cell lysates of indicated cell lines were separated on SDS PAGE and analysed by Western blotting using the indicated antibodies. (C) Indicated cell lines were treated with 1 μM MLN4924 in 0.1% DMSO, or just DMSO at 0.1% for 24 h. Samples were further processed as in (B). (D) Indicated cell lines were analysed on a Canto flow cytometer. Shown populations were preselected for cells and single cells before defining the gate for GFP positive cells (shown). GFP-A is plotted against PE-A in all cases. Numbers indicate percentage of cells within the respective gate. (E) Histogram representation of plots in (D). KI = A549 KRAS_GFP/GFP_ cells (referred to as a549_GFPKRAS_ cells throughout text). Western blots are representative of at least 3 independent experiments. Flow cytometry data are representative of 2 independent experiments.

### AdPROM mediated degradation of endogenous RAS proteins

The AdPROM-mediated degradation of GFP-K-RAS in A549_GFPKRAS_ cells demonstrated the prospect of targeted degradation of endogenous K-RAS. However, the presence of the GFP-tag raised the possibility of ubiquitination occurring on the GFP moiety, instead of on K-RAS. Therefore, we sought to explore whether we could exploit the AdPROM system to degrade endogenous, unmodified K-RAS from A549 cells. At present, there are no reported high affinity, selective polypeptide binders of K-RAS. However, we utilized an anti-H-RAS (aHRAS) monobody that was reported to bind and immunoprecipitate both H-RAS and K-RAS, but not N-RAS [36]. Using this monobody with a FLAG-tag, we showed that anti-FLAG immunoprecipitates (IPs) could robustly coprecipitate both GFP-tagged and untagged K-RAS, as well as the lower molecular weight protein representing the H- and/or N-RAS band but most likely to be H-RAS [36] (Fig. 3A). However, neither RAS protein was completely depleted from flow-through extracts, suggesting incomplete immunoprecipitation (Fig. 3A). In contrast, anti-FLAG IPs from extracts expressing Flag-VHL control did not co-precipitate either protein (Fig. 3A).

**Figure 3.**
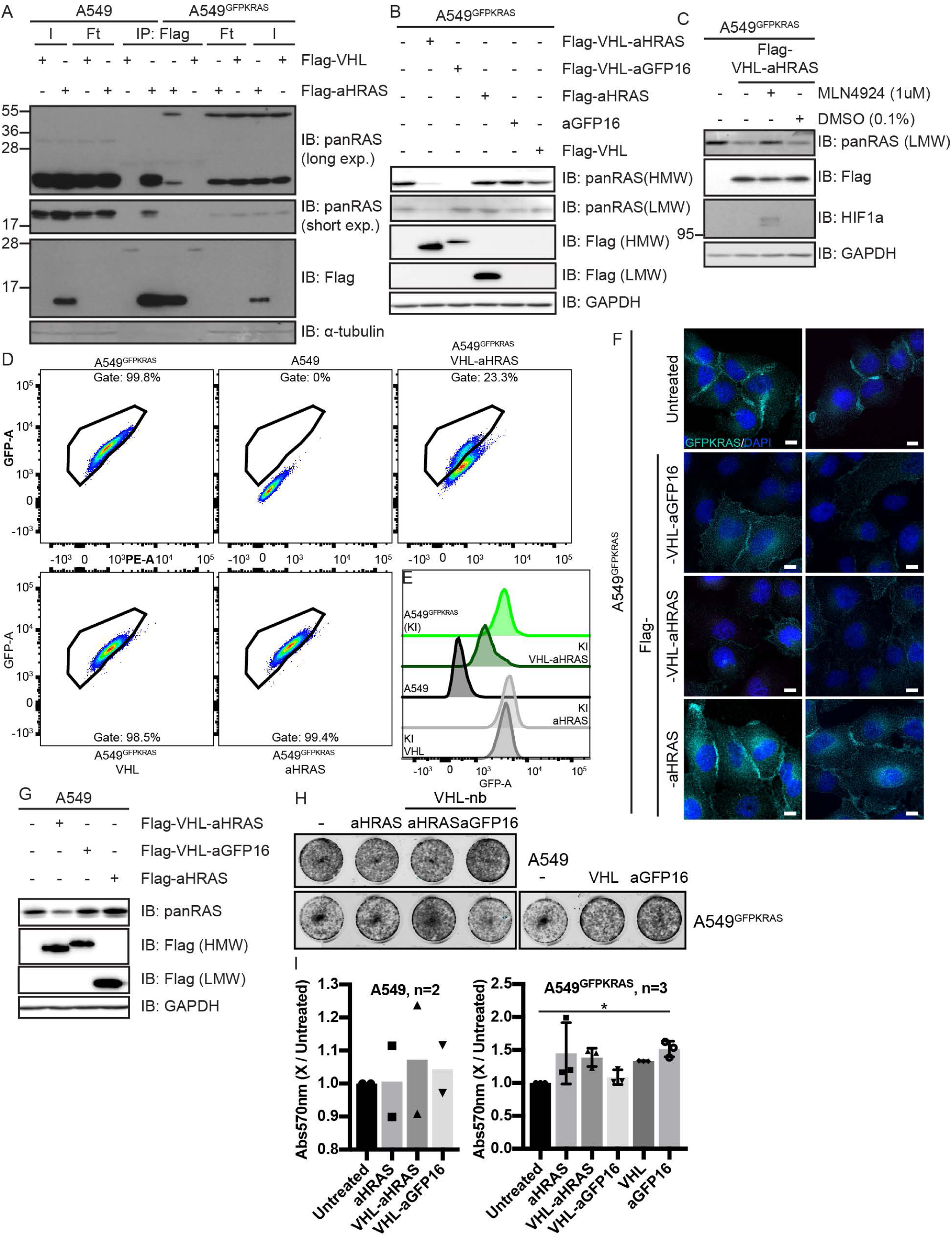
Degradation of endogenous RAS using a RAS-specific monobody. (A) Cell lysates of indicated cell lines were subjected to immunoprecipitation with anti-Flag beads. Input (I), Flowthrough (Ft) and precipitates (IP) were run on SDS-PAGE and subjected to Western blotting with the respective antibodies. (B), (C) and (G) SDS-PAGE and Western blots of lysates of indicated cell lines using the indicated antibodies. Samples were treated with 1 μM MLN4924 or 0.1% DMSO for 24 h (C). (D) and (E), flow cytometric analysis of indicated cells, done as in Figure 2. KI = A549_GFPKRAS_ cells. (F) Widefield immunofluorescence microscopy of indicated cell lines treated with anti-GFP antibody and DAPI for staining. Scalebar = 10 μm. Two representative images are shown for each condition. (H) 5,000 cells from (B) and (G) were grown in triplicate in 12 well dishes. After 7 days, cells were fixed and stained with crystal violet. A representative image of the replicates is shown. (I) Staining from plates in (H) was extracted by methanol and absorbance at 570 nm was measured. Plotted 570 nm values are relative to the respective untreated sample. The number of biological replicates is indicated next to the cell line name. For statistical analysis, one-way ANOVA analysis with Dunnett’s multiple comparisons test was done. Comparisons were drawn to the untreated sample. Western blots and immunofluorescence data are representative of at least 3 independent experiments. Flow cytometry data are representative of 2 independent experiments.

Next, we sought to investigate whether AdPROM consisting of VHL fused to aHRAS monobody could target K- and H-RAS proteins for degradation. In A549_GFPKRAS_ cells, the expression of VHL-aHRAS resulted in a strong reduction of the GFP-K-RAS protein levels when compared to untransduced, VHL or monobody alone controls (Fig. 3B). However, the degradation induced by VHL-aHRAS AdPROM was slightly less efficient than that achieved with the VHL-aGFP16 AdPROM (Fig. 3B). Unlike VHL-aGFP16, VHL-aHRAS also reduced the protein levels corresponding to the H-RAS and/or N-RAS band (Fig. 3B). The loss in protein levels of endogenous H-RAS protein caused by VHL-aHRAS AdPROM could be rescued by the Cullin neddylation inhibitor MLN4924, suggesting that the degradation was mediated through CUL2-RBX E3 ligase machinery (Fig. 3C). As expected, MLN4924 also stabilised endogenous HIF1α (Fig. 3C). We also assessed the relative abundance of GFP-K-RAS in mixed populations of A549_GFPKRAS_ cells transduced with VHL-aHRAS AdPROM in comparison to controls by flow cytometry. We found that 77% of cells showed degradation of GFP-K-RAS, as assessed by the shift of the GFP-positive gated population towards the GFP-negative population (Fig. 3D) and the overall reduction of GFP-signal (Fig. 3E). The remaining 23% of cells transduced with VHL-aHRAS were seemingly unaffected in both positioning in the GFP-positive gate (Fig. 3D), as well as GFP intensity (Fig. 3E). Transductions with VHL or aHRAS alone did not induce a noticeable shift of the GFP population or GFP signal intensity (Fig. 3D & E).

Uneven retroviral transduction of cells could result in unequal expression of the AdPROM constructs in different cells resulting in a mixed, divergent cell population, which may account for the apparent uneven degradation of GFP-K-RAS through VHL-aHRAS. When we analysed these A549_GFPKRAS_ mixed cell populations by immunofluorescence for GFP signal, in untransduced and aHRAS-transduced control cells, a predominant plasma membrane GFP-K-RAS signal was evident (Fig. 3F). Transduction of A549_GFPKRAS_ cells with either VHL-aHRAS or VHL-aGFP16 AdPROM produced a heterogenous population comprising cells with missing or severely attenuated GFP signal, and cells with intact GFP-K-RAS staining pattern, localizing mainly to the plasma membrane (Fig. 3F). In contrast, we noticed a slight increase in perinuclear GFP-K-RAS signal in cells transduced with the aHRAS monobody alone (Fig. 3F). Interestingly, we detected that the majority of the monobody itself was in the nucleus (Fig. S4), while we were unable to consistently detect signals for the AdPROM fusion proteins by anti-FLAG immunofluorescence (Fig. S4).

We also tested the degradation of endogenous K- and H-RAS in WT A549 cells with VHL-aHRAS AdPROM. The transduction of cells with VHL-aHRAS resulted in a substantial reduction in apparent levels of both K-RAS (upper band) and H-RAS (lower band) proteins as detected by the pan-RAS antibody compared to untransduced controls (Fig. 3G). Unlike in A549_GFPKRAS_ cells (Fig. 3B), WT cells transduced with VHL-aGFP16 AdPROM did not have any noticeable effect on K-RAS and H-RAS protein levels relative to untransduced cells (Fig. 3G), further validating the targeted nature of RAS degradation by AdPROM. Cells transduced with the aHRAS monobody alone led to a slight increase in abundance of both K-RAS and H-RAS proteins compared to untransduced controls (Fig. 3G). We sought to explore whether targeted degradation of K- and H-RAS proteins from WT A549 cells using the VHL-aHRAS AdPROM, and GFP-K-RAS from A549_GFPKRAS_ cells using the VHL-aGFP16 AdPROM would impact cell proliferation. No significant differences in proliferation could be observed for either WT A549 or A549_GFPKRAS_ cells following AdPROM-mediated degradation of the respective RAS proteins compared to controls after 7 days, as measured by crystal violet staining (Figs. 3H & I). Although A549 cells harbour the oncogenic K-RAS_G12S_ mutation, they also harbour over 250 genetic mutations (COSMIC cell lines project), including some known oncogenes and tumour suppressors reducing the likelihood that these cells are solely dependent on the K-RAS_G12S_ oncogene for their proliferation.

### Expansion of the RAS-targeting AdPROM system in different cell lines

Having demonstrated for the first time that the VHL-aHRAS AdPROM system could target endogenous H- and K-RAS for degradation in A549 cells, we sought to explore whether the system would work in other cell lines. First, we compared different cell lines for their endogenous RAS protein expression (Fig. 4A) relative to A549 cells. All cells tested displayed K-RAS protein expression similar to or slightly lower than A549 cells. SW620 cells, which harbour the G12V mutation on K-RAS [37], displayed similar levels of expression to A549 cells, however, we noticed that K-RAS in this cell line produced a slight but noticeable molecular weight shift, when probed with panRAS and K-RAS4B antibodies (Fig. 4A). Protein levels corresponding to the lower H- and/or N-RAS band were similar in all lines tested but overall much lower in intensity than that seen for K-RAS. We tested the ability of VHL-aHRAS AdPROM to degrade RAS proteins from HT-29 and SW620 cells. In HT-29 cells, which express WT RAS proteins but harbour the activating BRAF V600E mutation [38], only the levels of H-RAS but not K-RAS proteins were reduced by VHL-aHRAS AdPROM compared to controls (Fig. 4B, left panel). The proliferation of HT-29 cells was only reduced by about 50% by the aHRAS monobody alone (Fig. 4C and D), while the VHL-aHRAS and VHL-aGFP16 constructs reduced growth to a lesser extent (Fig. 4D, left panel). For SW620 cells, which harbour the G12V mutation of K-RAS, we noticed a high K-RAS signal to H-/N-RAS signal ratio, as the latter was barely detectable (Fig. 4B, right panel). We observed stabilization of K-RAS with the aHRAS monobody alone, while VHL-aHRAS failed to degrade K-RAS compared to controls. Interestingly, both the aHRAS monobody alone and the VHL-aHRAS AdPROM but not VHL-aGFP16 AdPROM were able to reduce the proliferation of SW620 cells significantly by about 50% (Fig. 4C & D).

**Figure 4.**
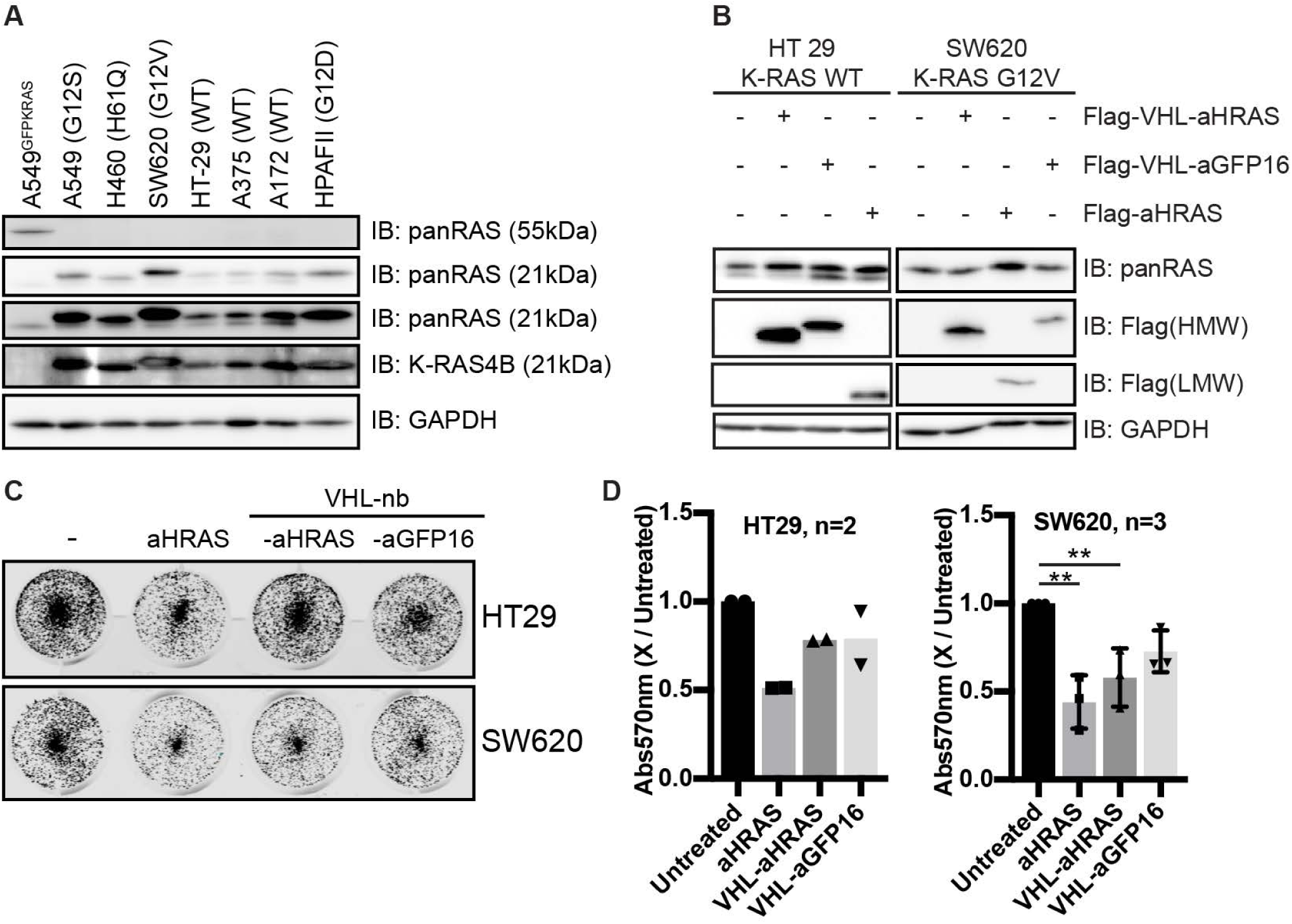
Degradation of RAS in different cell lines and effects on proliferation. Lysates of untreated (A), or retrovirally transduced cell lines (indicated expression constructs) (B) were separated by SDS PAGE and analysed by Western blotting with the indicated antibodies. Comparison of cell lines in (A) was done only once. K-RAS mutation statuses for individual cell lines are indicated in brackets. (C) 5,000 cells from (B) were grown in triplicate in 12-well dishes. After 7 days, cells were fixed and stained with crystal violet. A representative image of the replicates is shown. (D) Staining from plates in (C) was extracted by methanol and absorbance at 570 nm was measured. Plotted 570 nm values are relative to the respective untreated sample. The number of biological replicates (applies to Western blots in (B) as well) is indicated next to the cell line. For statistical analysis one-way ANOVA analysis with Dunnett’s multiple comparisons test was done. Comparisons were drawn to the untreated sample.

## Discussion

In this report, we demonstrate that endogenous K-RAS and H-RAS proteins can be targeted for degradation using the proteolytic AdPROM system. RAS proteins have remained elusive targets for anti-cancer therapies, primarily due to their undruggability [1]. Research into obtaining small molecule inhibitors of K-RAS has been carried out for over 30 years without much success [39]. Recently, RAS targeting small molecules have emerged, with specificities to (i) a specific mutation status of K-RAS (G12C), i.e. ARS-1620 [40], and ARS-853 [41]; (ii) K-RAS, independent of the mutation status [42]; or (iii) RAS proteins in either nucleotide binding state [43]. Two compounds targeting K-RAS_G12C_ mutation, AMG510 and MRTX849, are currently undergoing clinical trials [44]. An alternative approach has been the development of high affinity polypeptide binders of RAS that neutralise the RAS function. A class of binders based on ankyrin repeat proteins (DARPINs) [45] can bind and neutralise specific nucleotide loading states of RAS proteins [45]. Similarly, a fibronectin type III domain-based RAS-binding monobody [36,46–48] was shown to bind and inhibit the dimerization of both K- and H-RAS, and the overexpression of this monobody was shown to suppress tumour growth in mice [48]. Besides inhibition, RAS degradation offers another alternative approach at inhibiting RAS function to target RAS-dependent cancer cells. In this context, the dTAG-13 PROTAC was used to degrade FKBP12_F36V_-tagged K-RAS [17] through the UPS, albeit when overexpressed in cells. Our AdPROM system, demonstrating here that endogenous RAS proteins can be targeted for proteolysis through the UPS, informs that small molecules targeting RAS proteins for degradation is a viable option for intervention. Furthermore, our A549_GFPKRAS_ cells provide an excellent high throughput screening platform to test the efficacy of such molecules. However, targeted delivery of polypeptide binders of RAS proteins or the proteolytic AdPROM system into RAS-dependent cancer cells remains challenging and therefore currently offers limited therapeutic potential.

One difficulty in the study of RAS proteins is the absence of robust reagents to reliably detect specific RAS proteins at the endogenous levels, especially by immunofluorescence [32]. Often, overexpression of GFP-tagged or other epitope-tagged K-RAS has been employed to investigate RAS localization [36,49,50]. Therefore, our homozygous A549_GFPKRAS_ NSCLC cell line generated using CRISPR/Cas9, notwithstanding the potential caveats of GFP-tagging, has allowed us to not only assess localization of endogenously driven GFP-K-RAS protein but its mobility shift has allowed us to test the utility of panRAS and K-RAS antibodies in detecting K-RAS by Western blotting. Beyond the plasma membrane localisation, we observed additional disperse cytoplasmic signals of endogenous GFP-K-RAS, but no mitochondrial localisation. When overexpressed, K-RAS_G12V_ has been implied to be transported into mitochondria, leading to alterations of membrane potential, a decrease in respiration and an increase in glycolysis [51]. Potential compartments for the observed cytosolic signal for K-RAS could be Golgi, as seen for H- and N-RAS [52], which could correspond to K-RAS4A signal, or Endoplasmic Reticulum. However, this remains to be verified.

While the VHL-aGFP AdPROM was very effective at selectively degrading GFP-K-RAS from A549_GFPKRAS_ cells, the VHL-aHRAS AdPROM degraded endogenous H- and K-RAS with mixed efficacy in different cell lines. In developing the aHRAS monobody, the authors noted a difference in downstream behaviours of H- and K-RAS upon monobody binding, such as K-RAS, but not H-RAS being displaced from the membrane, or the mutant K-RAS, but not mutant H-RAS interaction with RAF being disturbed by monobody binding [46]. The full determinants of interaction between the aHRAS monobody and different H- and K-RAS mutants or any post-translationally modified forms remain poorly defined. It is perhaps the differences in affinity between the RAS proteins and the aHRAS monobody that define how robustly or poorly VHL-aHRAS can degrade different RAS proteins. Nonetheless, our study proves that any high-affinity polypeptide binders that can selectively bind specific RAS proteins or mutants can be packaged with VHL-AdPROM in order to target specific RAS proteins for proteasomal degradation. We also noted that aHRAS monobody alone resulted in a marked stabilization of both H-RAS and K-RAS in multiple cells (Fig. 3G & F and Fig. 4B), which could be caused either by a feedback loop induced by the inhibition of both RAS species imparted by aHRAS binding, or by blocking the natural turnover pathway through binding the RAS dimerization interface at helical structures α4-α5 [36].

For the cell lines that we used, AdPROM-mediated degradation of H-/K-RAS was not sufficient to induce inhibition of anchorage-dependent cell proliferation. For A549 cells that are considered not to be K-RAS-dependent for proliferation, this is perhaps not surprising [53, 54]. Meanwhile, SW620 cells have been reported to be K-RAS dependent for proliferation [55], however, their proliferation was inhibited by aHRAS monobody alone and the VHL-aHRAS AdPROM, which caused no detectable degradation of K-RAS, did not inhibit their proliferation any further. The inhibition of cell proliferation of RAS-dependent cells by aHRAS monobody is consistent with previous reports [36, 48]. The lack of degradation of K-RAS by VHL-aHRAS AdPROM could be due to the unusual size shift of K-RAS in these cells, possibly caused by a post-translational modification or a mutation that might allow binding to aHRAS monobody but prevent ubiquitylation by the VHL-AdPROM, although this needs to be defined further. Many RAS-dependent cell proliferation assays employ anchorage-independent 3D cultures. For example, the K-RAS_G12C_ drug ARS-1620 was shown to be effective at inhibiting RAS-dependent cell proliferation in 3D cultures but not in 2D cultures [40]. In order to assess the effects of AdPROM-mediated degradation of H-/K-RAS on proliferation robustly, it will be essential to first obtain polypeptide RAS binders that bind to specific RAS proteins with high affinity and then use them in RAS-dependent cell lines using 3D proliferation assays.

Recently two allosteric small molecule binders were described for K-RAS with low micromolar and nanomolar binding affinities [42, 43]. It would be important to test these binders’ capabilities as K-RAS targeting warheads in a PROTAC approach. In this line, a re-evaluation of RAS binding molecules, with or without inhibitory function, might prove successful for PROTAC designs.

## Conclusion

Our findings demonstrate clearly that endogenous RAS proteins can be targeted for proteasomal degradation by employing the AdPROM system. The system is not only suitable for studying the functions of these RAS proteins but also unequivocally informs that targeted proteolysis of endogenous K-RAS is a viable strategy to target K-RAS-dependent pathologies. The findings open up exciting opportunities to develop VHL-recruiting K-RAS-specific cell-permeable PROTACs as potential therapeutic agents. Our findings also highlight the need for developing better and more selective RAS binding polypeptides, such as nanobodies or monobodies, to achieve more selective degradation with the AdPROM system.

## List of abbreviations

AdPROM: Affinity directed PROtein Missile
ASC: Apoptosis-associated speck-like protein containing a CARD
ATPB: ATP synthase subunit β
BRAF – B: Rapidly Accelerated Fibrosarcoma
Cas9: CRISPR associated protein 9
CRBN: Cereblon
CRISPR: Clustered Regularly Interspaced short palindromic repeats
CUL: Cullin
DAPI: 4,6-diamidino-2-phenylindole
DARPIN: Designed Ankyrin Repeat Protein
FACS: Fluorescence Activated Cell Sorting
FKBP: FK506 Binding Protein
GAP: GTPase Activating Protein
GAPDH: Glyceraldehyde 3-phosphate dehydrogenase
GDP: Guanosine diphosphate
GEF: Guanosine Nucleotide Exchange Factor
GFP: Green Fluorescent Protein
GTP: Guanosine triphosphate
HIF1α: Hypoxia Inducible Factor 1α
HVR: Hypervariable region
MAPK: Mitogen Activated Protein Kinase
NSCLC: Non-small cell lung carcinoma
PAWS1: Protein associated with Smad1
PI3K: Phosphoinositide 3-kinase
qRT-PCR: Real-time quantitative RT-PCR
RAS: Rat Sarcoma
RBX1: RING box protein 1
SDS-PAGE: SDS Polyacrylamide Gel Electrophoresis
SHP2: Src homology region 2 (SH2)-containing protein tyrosine phosphatase 2
UPS: Ubiquitin Proteasome System
VHL: Von-Hippel-Lindau
VPS34: Vacuolar protein sorting 34

## Declarations

### Ethics approval and consent to participate

Not applicable

### Consent for publication

Not applicable

### Availability of data and materials

All data generated and analysed during this study is currently available in the Center for Open Science repository under the following link, https://osf.io/g5qn9/?view_only=4ef0cf7df11f4174b5f6760fa10042fe Data will be stored as permanent registry and publicly available upon acceptance of the manuscript.

### Authors’ contributions

TJM generated all used plasmids. SR, AK, MAQ and GPS designed the project. SR and GPS drafted the manuscript. SR acquired and analysed the data. SR and GPS interpreted the data.

## Acknowledgements

We thank GS laboratory members for their highly appreciated experimental advice and/or discussions during the course of these experiments. We thank L. Fin, E. Allen, J. Stark and A. Muir for help and assistance with tissue culture, the staff at the DNA Sequencing services (School of Life Sciences, University of Dundee), and the cloning teams within the MRC PPU reagents and services (University of Dundee), coordinated by J. Hastie and H. McLauchlan. We thank the staff at the Dundee Imaging Facility (School of Life Sciences, University of Dundee), and the staff at the flow cytometry facility (School of Life Sciences, University of Dundee) for their invaluable help and advice throughout this project.

---

**Figure S1.**
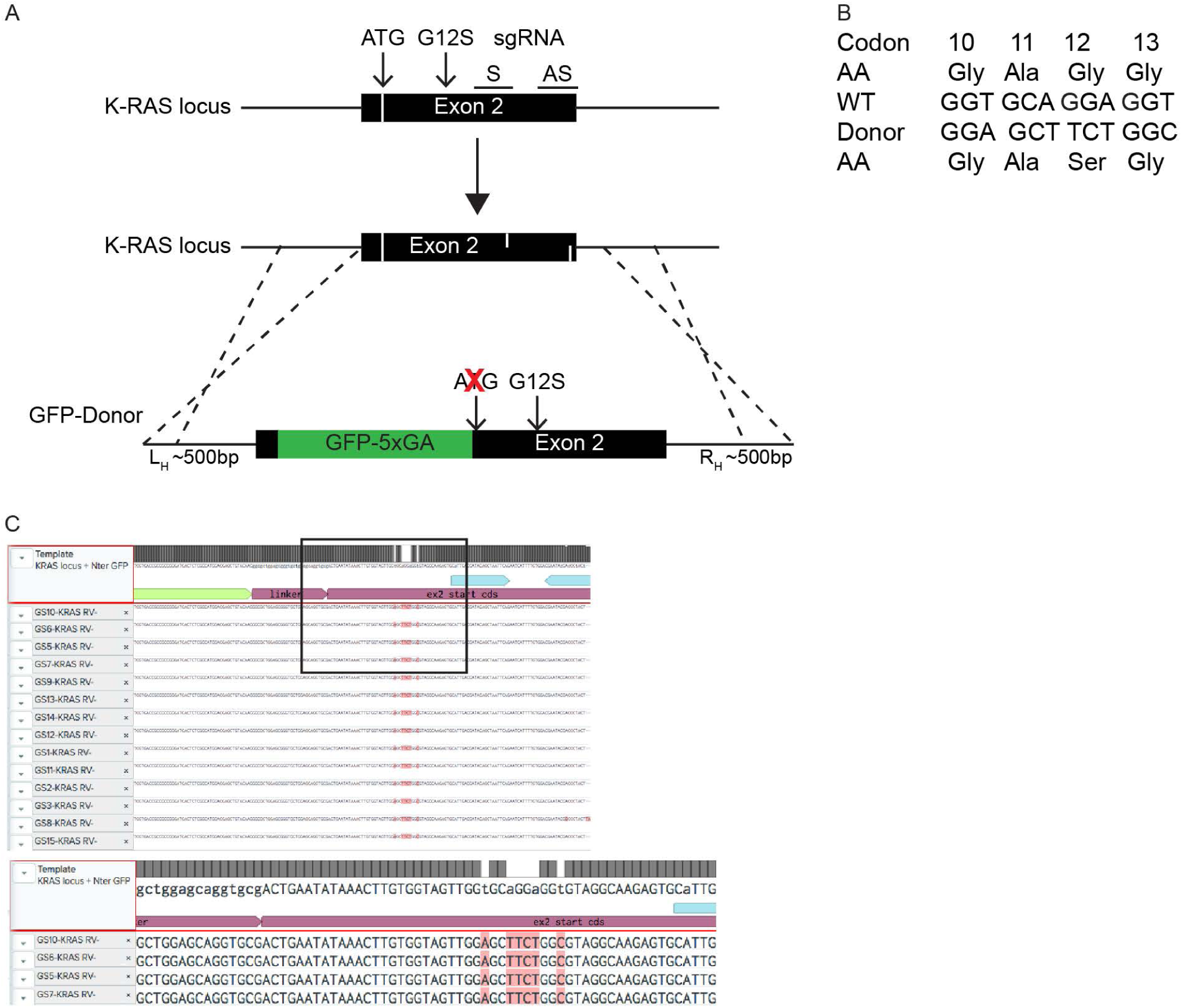
Characterization of A549^GFPKRAs^. (A) Schematic representation of the CRISPR/Cas9 strategy used for A549 cells. Two plasmids encoding sgRNA sequences targeting the K-RAS locus on exon 2 were co-expressed with Cas9-D10A, to create two nicks in K-RAS complementary strands for a double stranded break. A donor plasmid consisting of GFP cDNA sequence without the stop codon followed by a GAGAGAGAGA linker flanked by Left and Right homology arms (L_H_ and R_H_, respectively) was designed and co-transfected to allow homologous recombination for insertion of the GFP-5xGA tag onto the native K-RAS locus at the start codon. Consequently, the start codon of K-RAS was eliminated. (B) The indicated silent mutations on sgRNA target codons (10-13) were introduced in the donor sequence to block subsequent dsDNA breaks following integration of the donor on K-RAS locus. (C) A screenshot of DNA sequence analysis from Benchling of the resulting GFP-positive clone. Top: 14 DNA sequence files were aligned against the predicted WT RAS gene locus sequence with the GFP-fusion (indicated in light green) and the 5xGA linker. The box indicates area of the image that is magnified below. The magnified area shows DNA sequence alignment of the A549^GFPKRAs^ cell line at the site of the G12S mutation, which also shows silent mutations. The Ab1 files containing the DNA sequence chromatograms are deposited online.

**Figure S2.**
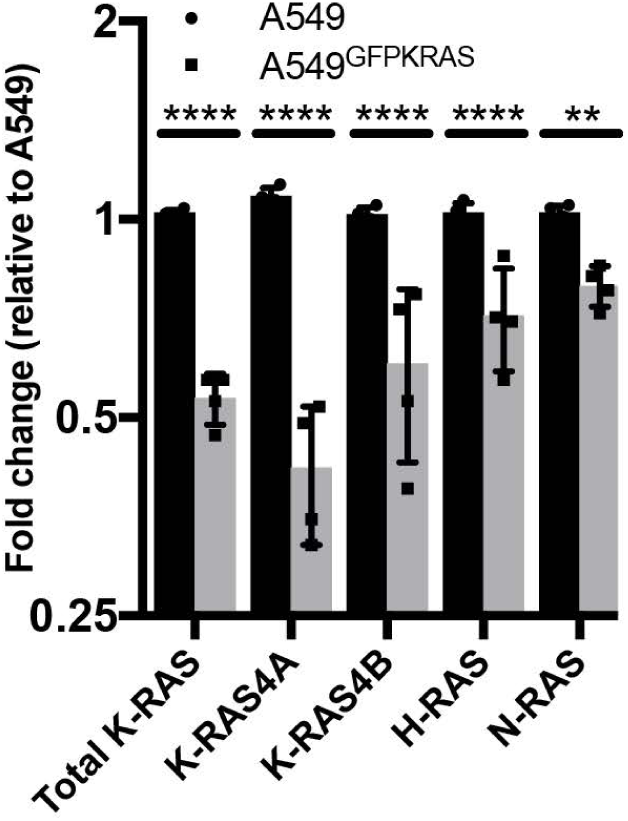
Analysis of RAS transcript levels in A549 WT and A549^GFPKRAs^. mRNA was extracted from indicated cell lines and cDNA was synthesised. Fold changes in RAS expression were calculated after qRT-PCR with specific primers between A549^GFPKRAs^ (grey bars) and A549 WT (black bars). Error bars are shown for n=4. Statistical significance was calculated with a 2-way ANOVA, Sidak’s multiple comparison test.

**Figure S3.**
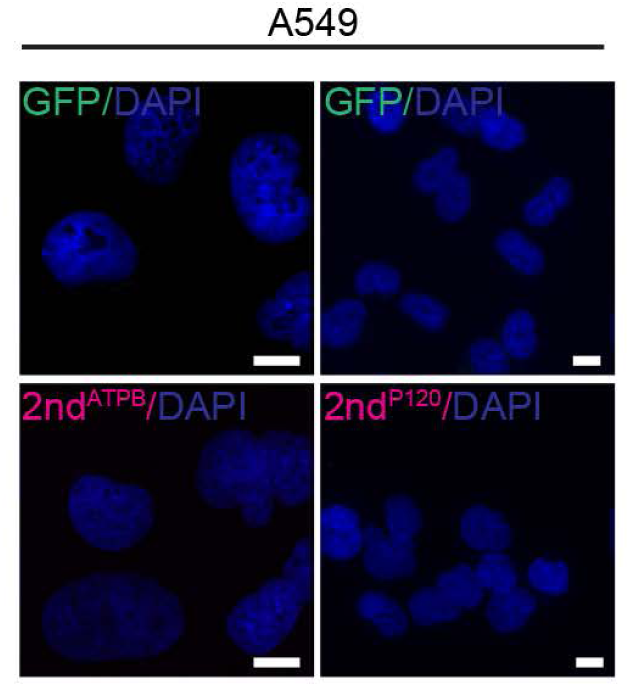
Negative controls for Figure 1D. Experimental procedure is described in Figure 1D. An additional slide of A549 WT cells (GFP negative) was treated with the secondary antibody used for ATPB or P120. Exposure times are set to be the same as for the positive sample slides in Figure 1D. Scalebar = 10µm

**Figure S4.**
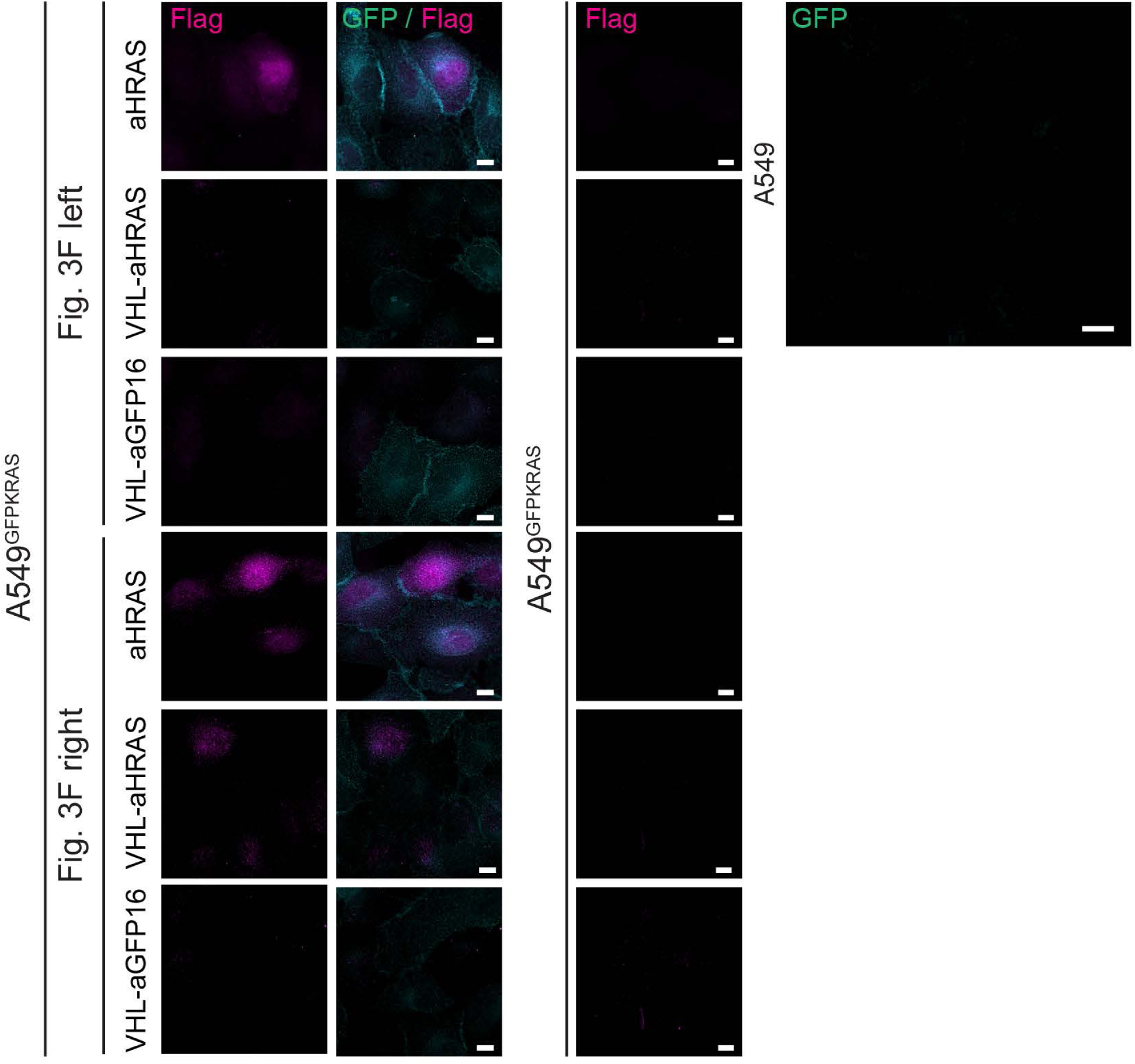
FLAG signal and controls for Figure 3F. Slides of Fig. 3F (A549^GFPKRAs^ with or without indicated transductions) were additionally stained with anti-FLAG antibody and secondary stain for detection in the 594 channel (Flag) (2 leftmost columns). The top 4 rows represent the left column of Fig. 3F, the bottom 4 rows the right column of Fig. 3F. Appropriate negative control (A549^GFPKRAs^ cells without a FLAG construct) were stained with the same antibody combination and exposure time (Column 3). Brightness and contrast for individual positive stains were background adjusted for these samples (note that the same negative sample is used for more than one picture). Additionally A549 WT cells were stained with the GFP antibody to act as negative control for GFP signal in main Figure 3F (panel on the right). Scalebar = 10µm.

